# Evidence for modulation of EEG microstate sequence by vigilance level

**DOI:** 10.1101/2020.02.26.966150

**Authors:** Marina Krylova, Sarah Alizadeh, Igor Izyurov, Vanessa Teckentrup, Catie Chang, Johan van der Meer, Michael Erb, Nils Kroemer, Thomas Koenig, Martin Walter, Hamidreza Jamalabadi

**Affiliations:** Department of Psychiatry and Psychotherapy, Division for Translational Psychiatry, University of Tübingen, Tübingen, Germany; Department of Electrical Engineering and Computer Science, Vanderbilt University, Nashville, USA; Clinical Affective Neuroimaging Laboratory, Magdeburg, Germany; Leibniz Institute for Neurobiology, Magdeburg, Germany; QIMR Berghofer Medial Research Institute, Brisbane, Australia; Division of Biomedical Magnetic Resonance, University of Tübingen, Tübingen, Germany; Max Planck Institute for biological cybernetics, Tübingen, Germany; Translational Research Center, University Hospital of Psychiatry and Psychotherapy, University of Bern, Switzerland; Department of Psychiatry and Psychotherapy, Jena University Hospital, Philosophenweg 3, 07743 Jena, Germany

**Author notes:** **Corresponding Authors:** Prof. Dr. Martin Walter Department of Psychiatry and Psychotherapy, Jena University Hospital, Philosophenweg 3, 07743 Jena, Germany; Dr. rer. nat. Hamidreza Jamalabadi, Department of Psychiatry and Psychotherapy, Division for Translational Psychiatry, University of Tübingen, Calwerstr. 14, 72076 Tübingen, Germany. Highlights - EEG microstate parameters are strongly related to vigilance levels and can predict them - We find that vigilance Granger-causes changes in parameters of microstates - Duration and occurrence of EEG microstates are differentially modulated by vigilance level.

**Keywords:** EEG microstates, vigilance, global signal

## Abstract

The momentary global functional state of the brain is reflected in its electric field configuration and cluster analytical approaches have consistently shown four configurations, referred to as EEG microstate classes A to D. Changes in microstate parameters are associated with a number of neuropsychiatric disorders, task performance, and mental state establishing their relevance for cognition. However, the common practice to use eye-closed resting state data to assess the temporal dynamics of microstate parameters might induce systematic confounds related to vigilance levels. Here, we studied the dynamics of microstate parameters in two independent data sets and showed that the parameters of microstates are strongly associated with vigilance level assessed both by EEG power analysis and fMRI global signal. We found that the duration and contribution of microstate class C, as well as transition probabilities towards microstate class C were positively associated with vigilance, whereas the sign was reversed for microstate classes A and B. Furthermore, in looking for the origins of the correspondence between microstates and vigilance level, we found Granger-causal effects of vigilance levels on microstate sequence parameters. Collectively, our findings suggest that duration and occurrence of microstates have a different origin and possibly reflect different physiological processes. Finally, our findings indicate the need for taking vigilance levels into consideration in resting-sate EEG investigations.

## Introduction

The topographical distribution of brain electrical potentials reflects the large-scale brain activity which can be effectively measured using multichannel scalp EEG (Fallgatter et al 1997, Fallgatter et al 2001, Hallez et al 2007, Lehmann & Skrandies 1984). Intriguingly, these topographical configurations do not change randomly but remain quasi-stable for a short period of time of around 80 milliseconds before rapidly switching to another quasi-stable topography. These reoccurring stable geometrical patterns are referred to as EEG microstates (Lehmann et al 1987). Clustering approaches have consistently revealed four prototypical topographies which are sufficient to explain around 80% of variance in resting-state recordings (Khanna et al 2014, Khanna et al 2015, Koenig et al 2002, Michel & Koenig 2017), where the polarity can invert reflecting oscillations of the dominant generators.

As expected, the temporal characteristics of EEG microstate sequences change in response to a large number of external and internal stimulations. These include but are not limited to content of spontaneous thoughts (Lehmann et al 2010), behavioral (Dimitriadis et al 2015, Milz et al 2016, Seitzman et al 2017) and global brain state (Faber et al 2005, Katayama et al 2007) as well as pharmacological manipulations (Schiller et al 2019). These characteristics are different in patients suffering from neuropsychiatric disorders, such as schizophrenia, depression, dementia and multiple others (for a review seeKhanna et al 2015, Michel & Koenig 2017).

An important but often neglected aspect of these works, however, is that most of these studies were conducted using eyes-closed resting state EEG recordings alone. During typical eyes-closed rest, subjects tend to sequentially transit from complete wakefulness towards drowsiness. Tagliazucchi and Lafus in 2014 showed that likelihood of subjects falling asleep during eyes-closed recordings is high, and approximately half of the participants loose wakefulness after 10 minutes (Tagliazucchi & Laufs 2014). Transition from complete wakefulness to sleep onset is characterized by strong occipital alpha power increase immediately after closing the eyes that is followed up by anteriorization of alpha power focus (De Gennaro et al 2005) and subsequent increase of delta and theta activity indicating further transition to drowsiness (Olbrich et al 2009, Strijkstra et al 2003). As classification of EEG microstate classes is strongly dependent on the power of alpha frequency (Lehmann et al 1987), it is reasonable to hypothesize that the temporal characteristics of EEG microstates might be affected by vigilance changes. Since patients suffering from neuropsychiatric disorders are known to have altered vigilance regulation pattern (Hegerl & Hensch 2014, Olbrich et al 2012, Strauss et al 2015), this hypothesis, should it get confirmed, can have strong implications in interpreting EEG microstate alterations in a clinical context (for a related topic see Zanesco et al 2020).

To test this hypothesis, we investigated the relation of the characteristics of EEG microstate parameters with vigilance levels using two independent data sets of simultaneous eyes-closed EEG/fMRI resting state recordings. These datasets allowed us to 1) test the association of EEG microstate parameters with vigilance estimates based on EEG as well as fMRI metrics, and 2) assess potential causal relationship between vigilance loss and changes in temporal dynamics of EEG microstate characteristics.

## 1. Methods

### 1.1 Data Acquisition

The analysis was performed on two independent data sets of simultaneous EEG/fMRI recordings. The first study is registered at ClinicalTrials.gov, number NCT02602275 (date of registration: 28/10/2015) and approved by the ethics committee of the University of Magdeburg as well as the Competent Authority (Federal Institute for Drugs and Medical Devices). The second study was approved by the Ethics Committee of the University of Tübingen, Germany. A written informed consent was signed by each participant prior to any study participation.

#### Data set 1

The first data set includes 12-min eyes-closed resting-state recordings of 39 healthy male volunteers (mean age 43.7 ± 9.8) over one session of simultaneous EEG and 3-Tesla fMRI. EEG data were acquired using the BrainAmp MR system (Brain Products) with a 64-channel EasyCap. One channel placed on the back was used for ECG detection. AFz was used as reference electrode and FCz as ground electrode. The sampling rate was 5000 Hz. To increase the quality of EEG in simultaneous EEG-fMRI recordings, EEG cap was augmented with six carbon wire loops (CWLs) (van der Meer et al 2016). Four CWLs were placed on the outer surface of the EEG cap at the left and right frontal and left and right posterior locations, and two CWLs were attached to the cables connecting the EEG cap to the EEG amplifier (BrainAmpMR Plus). Imaging data were acquired on a 3 Tesla Philips whole body MRI system (Philips Medical Systems, Hamburg, Germany). First, structural T1-weighted images for spatial normalization were measured using a turbo field echo (TFE) sequence (274 sagittal slices covering the whole brain, flip angle = 8°, 256 × 256 matrix, voxel size = 2.5 × 2.5 × 3 mm^3^). Whole brain BOLD resting-state data were acquired over 34 axial slices using an echo planar imaging (Randerath et al) sequence (TR = 2,000 ms, TE = 30 ms, flip angle = 90°, 96 × 94 matrix, field of view = 24 cm, voxel size = 2.5 × 2.5 × 3 mm^3^).

#### Data set 2

The second data set includes 10-min eyes-closed resting-state recordings of 20 healthy male volunteers (mean age 26.8 ± 7.6) over one session of simultaneous EEG and 3-Tesla fMRI. EEG data were acquired with the same parameters as in data set 1. Imaging data were acquired on a 3 Tesla Siemens Prisma whole body MRI system (Siemens Medical Solutions, Erlangen, Germany). First, structural T1-weighted images for spatial normalization were measured using a three-dimensional magnetization-prepared rapid gradient echo (MP-RAGE) sequence (192 sagittal slices covering the whole brain, flip angle = 9°, 256 × 256 matrix, voxel size = 1 × 1 × 1 mm^3^, PE-GRAPPA factor 2). Whole brain BOLD resting-state data were acquired over 30 axial slices using an echo planar imaging (Randerath et al) sequence (TR = 1,800 ms, TE = 35 ms, flip angle = 79°, 64 × 64 matrix, field of view = 19.2 cm, voxel size = 3 × 3 × 4 mm^3^).

### 1.2 Data Preprocessing

#### Electroencephalography

First, gradient artifacts were removed from the EEG data by a motion informed template subtraction realized by the Bergen EEG-fMRI toolbox (Moosmann et al 2009) using a MRI template waveform obtained from 25 MRI artifacts in a sliding window manner (Allen et al 2000). Next, EEG data was first bandpass filtered between 0.3Hz to 200Hz and down-sampled to 1000Hz. The helium pump and ballisto-cardiac (BCG) artifacts were then removed using the Carbon-wire loop technique (van der Meer et al 2016). Next, the data were segmented into 2s and 1.8s epochs for the data set 1 and 2 respectively (i.e. equivalent to the TR of the BOLD resting-state scans), and the epochs containing muscle and head movement artifacts (outliers in spectral power between 110 and 140 Hz) were removed. The channels that contained more than 50% of epochs with artifacts were interpolated using routines provided by EEGLAB (Delorme & Makeig 2004). Finally, ICA decomposition of the EEG data was performed and components reflecting eye movements, continuous muscle activity and residual MRI-artefacts were removed. Six subjects from data set 1 and one subject from data set 2 were excluded from further analysis because of their low EEG quality or technical problems during data acquisition.

#### Functional MRI

fMRI data were preprocessed using SPM12 (FIL, Wellcome Trust Centre for Human Neuroimaging, UCL, London, UK) toolbox. The first three volumes of each recording were excluded from the analysis. The functional corrections included slice time correction and realignment to the first image. The structural T1-weighted volume was registered to the mean functional image and segmented, in order to normalize functional and structural images to the Montreal Neurological Institute (MNI) template brain. Finally, normalized functional volumes were smoothed with a three-dimensional Gaussian kernel of 6 mm full-width-half-maximum. Global signal time course was then estimated by averaging the z-scored time-series across all voxels with a gray matter tissue probability of at least 60% (based on tissue probability maps from the SPM12 toolbox). Finally, the global signal time-series were smoothed by calculating the mean values within a non-overlapping window of 3 TR.

#### Microstate extraction

EEG microstate analysis was performed separately for each dataset using the EEGLAB plugin for Microstates version 1.1, developed by Thomas König (http://www.thomaskoenig.ch/index.php/software/microstates-in-eeglab). Artefact-free EEG data were re-referenced to average reference, bandpass filtered between 2 and 20 Hz and further down-sampled to 250 Hz. The Global Field Power (GFP) was calculated as the root of the mean of the squared potential differences at all electrodes from the mean of instantaneous potentials across electrodes. Since the topography remains stable around peaks of the GFP, they are the best representative of the momentary map topography in terms of signal to-noise ratio (Koenig et al 2005). All maps marked as GFP peaks were extracted and submitted to a modified k-means clustering algorithm to deduce four classes of map topographies (microstates) that maximally explain the variance of the GFP peak map topographies. These four classes of map topographies were then submitted to a full permutation procedure (Koenig et al 1999) to compute mean classes across participants. Using the mean microstate classes across subjects as templates, for all participants the EEG topographies at the moments of GFP peaks were assigned to one of these four microstates based on maximal Pearson correlation. Successive GFP peak maps assigned to the same class were recognized as belonging to one microstate. Time points between GFP peaks were assigned to the microstate class of the temporally closest GFP peak (Figure 1A). In each epoch the time points before the first and after the last detected GFP peak were excluded because microstates cannot be determined in these points. For each microstate class and each epoch, four parameters were estimated i.e. duration (i.e. mean time spent in the current MS class), occurrence (i.e. frequency of appearance of the current MS class), contribution (i.e. percentage of total time of recording spent in the current MS class), and transition probability. As the observed transition probability might be affected by differences in occurrence between microstate classes, we also estimated a random transition probability model as described by Lehmann et al. (Lehmann, Faber et al. 2005). The resulting transition probability was calculated as the difference between the observed transition probability and the one estimated from the random transition probability model. To get rid of the fast fluctuations, the time course of the microstate parameters was smoothed by calculating the mean value within a non-overlapping window of 3 TR (resulting in 6 s and 5.6 s window length for data set 1 and 2 respectively).

**Figure 1:**
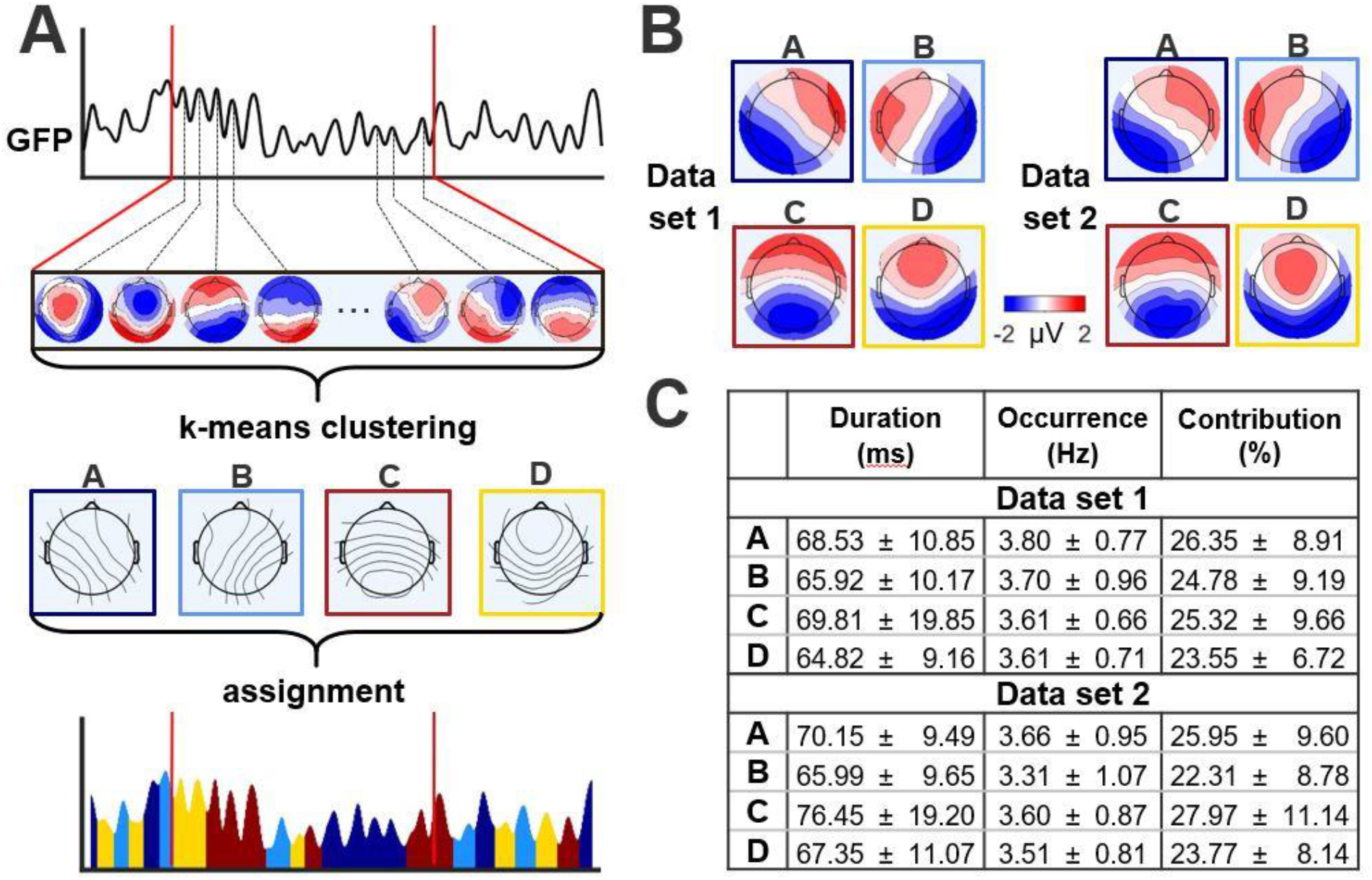
Microstate extraction and the parameters (A) Schematic representation of the microstate extraction procedure: 1) Global Field Power (GFP) is calculated at each time point of the multichannel EEG recording. 2) The head-surface potential maps at the peaks of the GFP curve are extracted and submitted to the clustering algorithm to reveal the dominant topographies (EEG microstates). 3) The original maps at peaks of the GFP curve are assigned to one of the microstate classes A, B, C, or D based on the degree of the spatial similarity with the microstate maps. (B) Head-surface topographies of the four EEG microstate classes for data set 1 (on the left) and data set 2 (on the right) during eyes closed resting. C: The microstate parameters for the four microstate classes (mean ± standard deviation).

### 1.3 Estimation of vigilance level

To obtain the temporal dynamics of vigilance fluctuation, power spectral density of EEG was estimated for each channel using a sliding Hamming window (Data set 1: 1457 points, 5.8s temporal width, and 65.7% overlap. Data set 2: 1311 points, 5.2s temporal width, and 65.7% overlap). The temporal resolution of the spectrogram was equivalent to the TR of fMRI resting state scans (i.e. 2s and 1.8s for the data set 1 and 2 respectively). Next, we estimated the global spectrogram by computing the root mean square (rms) value across all channels at each frequency. Then, vigilance time-series were calculated as rms amplitude in the alpha frequency band (7-13 Hz) divided by the rms amplitudes in the delta and theta frequency band (1-7 Hz) at each time point. The similar approach was used by (Falahpour et al 2018). To omit the fast fluctuations, the vigilance time-series were smoothed by calculating the mean value within a non-overlapping window of 3 TR.

## 2. Results

### 2.1 EEG microstates

Four EEG microstate classes (Figure 1B) explained on average 77.8 ± 2.9% and 76. 1 ± 3.6% of the total topographic variance across participants for data sets 1 and 2 respectively. Figure 1C shows mean duration, mean occurrence, and mean contribution for two datasets. The parameters of microstates are well in line with ranges reported in the literature (Khanna et al 2015, Kikuchi et al 2011, Kindler et al 2011, Milz et al 2016).

### 2.2 Correlation between vigilance level and microstate parameters

We analyzed the association between microstate and vigilance on two levels. First, the correlation between vigilance level and microstate parameters were calculated by estimating the Pearson correlation between mean vigilance and mean microstate parameters (see Figure 2). Mean vigilance and microstate parameters were estimated by averaging the time-series across all time points on the single subject level. We observed significant positive association of mean vigilance level with mean duration and contribution of microstate class C. Also, there was a negative association of mean vigilance level with occurrence and contribution of the microstate class A and B (see Appendix Table A.1 for details). Consistent with these findings, we also found a significant association of probability of transitions towards microstate class C from microstates class A but also microstate class D (see Appendix Table .2 for details). Interestingly, the higher vigilance level was characterized by increased duration of the microstate class C and decreased occurrence of the microstate classes A and B. Also, the pathway of the transitions between different microstate classes was altered together with changes in vigilance level, with higher probability for transitions towards microstate class C.

**Figure 2:**
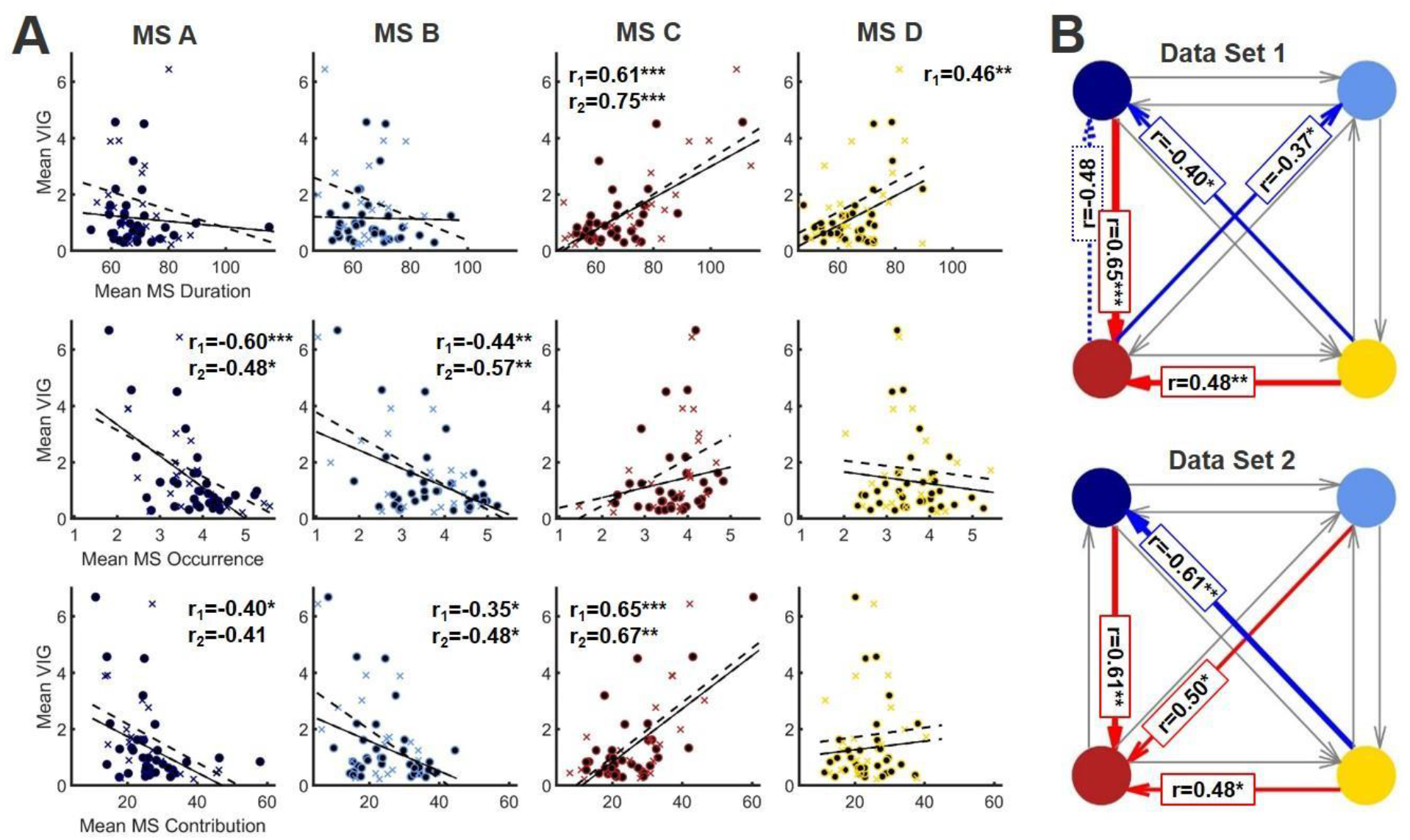
Association of the mean vigilance level and mean values of the EEG microstate parameters of the four microstate classes during eyes closed resting for the two data sets. A: Top panel: duration, middle panel: occurrence, bottom panel: contribution. Each dot represents data of one subject. Data for the first dataset are presented in circles and solid lines, data for the second dataset are presented in crosses and dashed lines. Mean duration and contribution of the microstate class C are positively, while occurrence and contribution of the microstates class A and B were negatively associated with vigilance level in both investigated datasets. B: The transition probabilities of the transitions between the four microstate classes. Red arrows represent positive association with vigilance level, blue – negative. The thickness of the line and the number of the stars corresponds to the p-value (p < 0.001 – thick / ***, 0.001 < p < 0.01 – medium / **, 0.01 < p < 0.05 – thin / *, trend level (p < 0.065) - dashed). For both data sets mean transition probabilities towards microstate class C were positively, while transition probabilities from microstate class D towards microstate class A were negatively associated with vigilance levels.

Second, we looked into temporal correlations between microstate parameters and vigilance level. For each subject the correlation between the vigilance time-series and the time courses of each of the microstate parameters were calculated. We used one-sample t-tests on the Fisher z-transformed correlation coefficients to estimate the group level effects. In line with the findings in the previous section, we observed significant positive association of the vigilance time course and time courses of the duration and contribution of the microstate class C. The time courses of the occurrence and contribution of the microstate class A and B were negatively associated with vigilance (see Appendix Table B.1 for details). However, we also observed (Figure 3 A), that the time course of the duration of the microstate class D was positively associated with vigilance time-series. Also, we observed a positive correlation between the vigilance time course and the time courses of the transition probability for the transitions from the microstate class A towards microstate class C and from the microstate class B towards microstate class D. The time course for of the probability of transitions from microstate class C towards microstate class B was negatively associated with the vigilance time-series (see Appendix Table B.2 for details).

**Figure 3.**
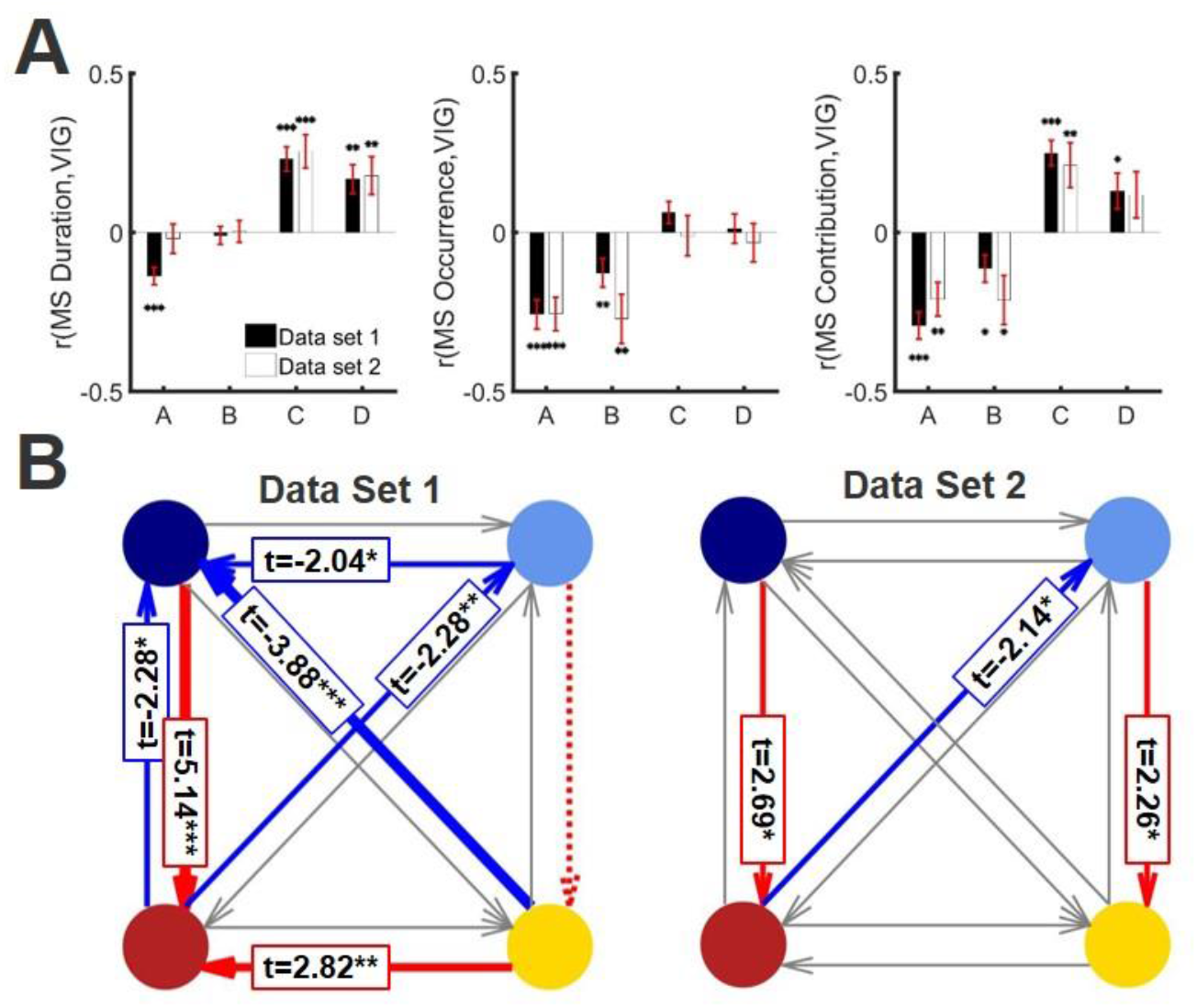
Association between the vigilance time-series and the time courses of the EEG microstate parameters during eyes closed resting for the two data sets. A: Top panel: duration, middle panel: occurrence, bottom panel: contribution. Bars show group average Fisher z-transformed correlation coefficients for the first (black) and second dataset (white), error bars represent standard deviation of the mean. Stars correspond to the significance levels (p < 0.001 - ***, 0.001 < p < 0.01 – **, 0.01 < p < 0.05 – *). Time courses of the duration and contribution of the microstate class C as well as duration of the microstate class D were positively, while time courses of the occurrence and contribution of the microstates class A and B were negatively associated with change of vigilance level in both investigated data sets. B: The transition probabilities of the transitions between the four microstate classes. Red arrows represent positive association, blue – negative. The thickness of the line corresponds to the p-value (p < 0.001 – thick, 0.001 < p < 0.01 – medium, 0.01 < p < 0.05 – thin, trend level (p < 0.065) - dashed). For both data sets time courses of the transition probabilities for transitions from microstate class A towards microstate class C as well as transitions from microstate class B towards microstate class D were positively, while the time course for of the probability of transitions from microstate class C towards microstate class B was negatively associated with vigilance time-series.

### 2.3 Multivariate pattern classification of the full vigilance time-series based on the microstate parameters

To test if the univariate correlations in section 3.2, allow for a temporal reconstruction of the vigilance level, we used support vector machine regression to predict the vigilance time-series based on the parameters of the microstates. We used 2-fold cross-validation on the subject level and repeated the cross-validation 100 times. To test against the null distribution, we randomly swapped microstate parameter time courses across different subjects and repeated the prediction procedure 1000 times. For both data sets, correlation between estimated and measures vigilance time-series were significantly above chance level (Figure 4).

**Figure 4:**
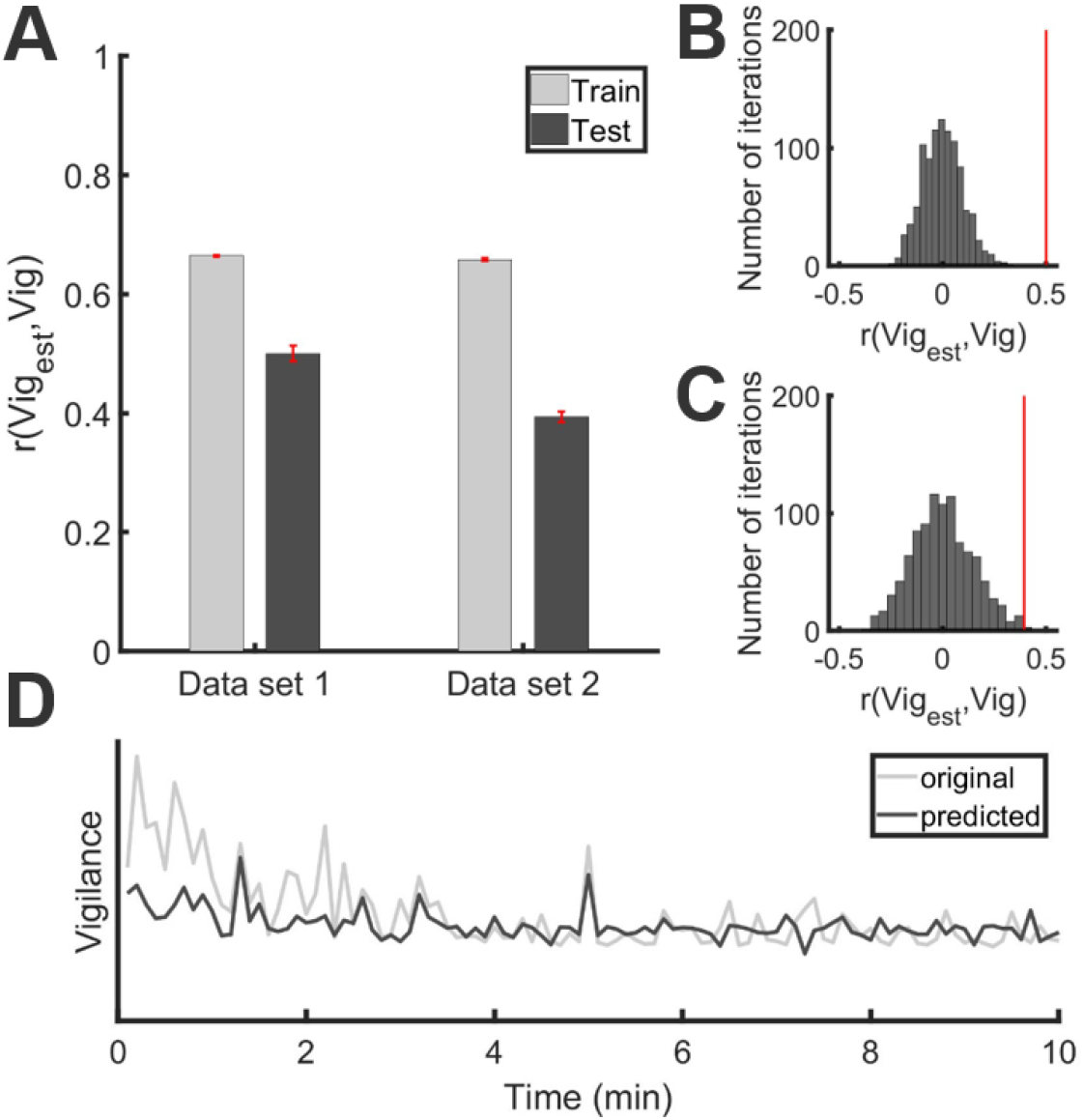
Prediction of the vigilance time-series based on the EEG microstate parameters using Support Vector Regression (SVR). A: correlation between estimated and real vigilance time-series for train (light gray) and test (dark gray). Error bars represent the standard deviation of the mean over 100 repetitions of the train and test procedure. B: null distribution for the first dataset. Vertical red line represents the mean test correlation between estimated and real vigilance time-series. C: null distribution for the second dataset. Vertical red line represents the mean test correlation between estimated and real vigilance time-series. D: example of the vigilance time-series prediction for a 10 min interval. Original data are shown in light gray and predicted in dark gray.

### 2.4 Correlation between time course of microstate parameters and BOLD global signal

Vigilance levels are known to be associated with fMRI global signal (Falahpour et al 2018, Wong et al 2016, Wong et al 2013). Here, we investigated the association of the temporal dynamics of the microstate parameters with the global signal. To do this, vigilance and microstate parameters’ time-series were convolved with the canonical hemodynamic response function to account for the hemodynamic delay. For each subject the correlation between the vigilance time-series and the time course of the fMRI global signal as well as the correlation between each of the microstate parameters and the time course of the fMRI global signal was calculated. We used one-sample t-tests on the Fisher z-transformed correlation coefficients to estimate the group level association.

We observed a significant negative correlation between the time course of vigilance levels and the time course of the global signal for both datasets (Figure 5 B). Additionally, we found a significant correlation between the duration and contribution of the microstate class C and contribution of microstate class A with the global signal (see Figure 5 A and Appendix Table C.1 for details). Interestingly, parameters of microstate class B did not show associations with the global signal. Along the same line, associations between the global signal time course and the time course of transition probabilities of microstates were weak and inconsistent across the two datasets (Figure 5 C and Appendix Table C.2 for details).

**Figure 5:**
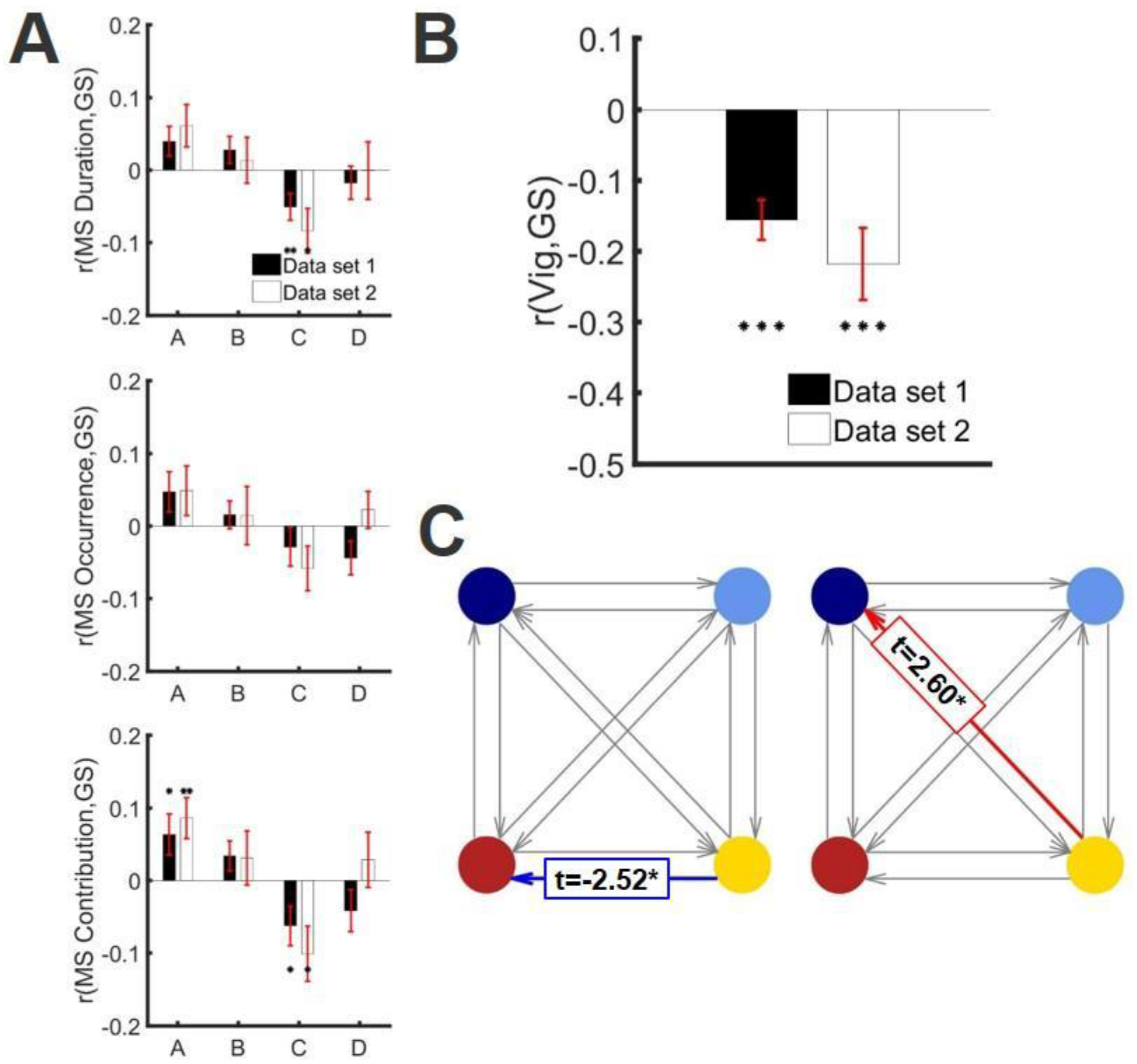
Association between the fMRI global signal time-series and the time course of the EEG microstate parameters during eyes closed resting for the two datasets. A: Top panel: duration, middle panel: occurrence, bottom panel: contribution. Bars show group average Fisher z-transformed correlation coefficients for the first (black) and second dataset (white), error bars represent standard deviation of the mean. Stars correspond to the significance levels (p < 0.001 - ***, 0.001 < p < 0.01 – **, 0.01 < p < 0.05 – *). Time courses of the duration and contribution of the microstate class C were negatively, while time course of the contribution of the microstate class A were positively associated with time course of global signal in both datasets. B: Association of the vigilance time-series and the global signal time-series for the for the first (black) and second dataset (white). Bars show group average Fisher z-transformed correlation coefficients, error bars represent the standard deviation of the mean. Stars correspond to the significance levels (p < 0.001 - ***, 0.001 < p < 0.01 – **, 0.01 < p < 0.05 – *). Time courses of the vigilance and global signal were negatively correlated in both datasets. C: The transition probabilities of the transitions between the four microstate classes. Red arrows represent positive association, blue – negative. The thickness of the line corresponds to the p-value (p < 0.001 – thick, 0.001 < p < 0.01 – medium, 0.01 < p < 0.05 – thin).

### 2.5 Causal effects of vigilance on microstate parameters

To estimate the temporal dynamics of the interplay between microstate parameters, vigilance fluctuations and global signal fluctuations, we calculated Granger causality (GC) which is a well-established measure of lag-based predictive causality. To this end, we used the MVGC toolbox (Barnett & Seth 2014). We calculated GC between time courses of the microstate parameters and vigilance time-series. For each participant within the two datasets, for each microstate class and each microstate parameter, we first estimated the optimal lag based on the Bayesian information criterion (BIC). As for most of the subjects (85.6% and 85.9% for the first and second dataset, respectively) the optimal model order (lag steps) was 1, we used this value for all other subjects. We then calculated time-domain pairwise-conditional GC values with a model order of 1 using the source and target pairs microstate parameter/vigilance. To estimate directionality, we obtained delta GC values, contrasting direction and modality. Hence, to estimate the extent to which the duration-based time-series of microstate A Granger-causes the vigilance time-series, GC values for the direction *Vig* → *MSA*_*duration*_ were subtracted from GC values for the direction *MSA*_*duration*_ → *Vig*. For statistical analyses, we bootstrapped the distribution of mean GC values with 50000 repetitions using bootci in Matlab2018a. We then calculated p-values by summing cases for which the bootstrapped mean GC value, depending on the direction of the effect, exceeded or went below zero and divided the sum by the number of iterations. Finally, to obtain two-tailed p-values we multiplied these values by two. We found that changes in vigilance cause changes in the temporal characteristics of microstate parameters (for details see Table 1).

**Table 1:** This table lists the results (two-tailed p-values of the bootstrapped statistics) of the causal relationships between time courses of the microstate parameters and vigilance time-series. Significant results (p < 0.05) are marked in bold. Stars correspond to the significance levels (p < 0.001 - ***, 0.001 < p < 0.01 – **, 0.01 < p < 0.05 – *). Changes in vigilance levels had causal effect on the changes of the parameters of microstates.

|  | Duration (ms) | Occurrence (Hz) | Contribution (%) |
| --- | --- | --- | --- |
| **Data set 1** |  |  |  |
| (Vig → A) – (A → Vig) | **p = 0.016\*** | **p < 0.001\*\*\*** | **p = 0.006\*\*** |
| (Vig → B) – (B → Vig) | p = 0.516 | p = 0.093 | p = 0.151 |
| (Vig → C) – (C → Vig) | p = 0.061 | p = 0.305 | p = 0.420 |
| (Vig → D) – (D → Vig) | **p < 0.001\*\*\*** | **p = 0.005\*\*** | p = 0.102 |
| **Data set 2** |  |  |  |
| (Vig → A) – (A → Vig) | p = 0.054 | **p < 0.001\*\*\*** | **p = 0.024\*** |
| (Vig → B) – (B → Vig) | p = 0.056 | **p = 0.002\*\*** | **p = 0.003\*\*** |
| (Vig → C) – (C → Vig) | p = 0.052 | **p = 0.026\*** | **p = 0.019\*** |
| (Vig → D) – (D → Vig) | **p = 0.018\*** | p = 0.226 | p = 0.892 |

## 3. Discussion

The temporal dynamics of microstates has been shown to reflect many cognitive processes and to be associated with a large number of psychiatric disorders. Here, looking into two independent datasets, we found that the temporal structure of microstates covaries with the vigilance level as measured using EEG frequency power but also fMRI based global signal. Importantly, we found evidence that microstates and vigilance levels are causally related and the changes in vigilance cause changes in the parameters of microstates. The observed associations had predictive power, and temporal dynamics of vigilance could be, to some extent, reconstructed based on the microstate parameters. The parameters of EEG microstates were highly associated with vigilance and global signal. In particular, we consistently found a relation between the duration of microstate C to both vigilance levels and global signal. We found as well that occurrence but not the duration of microstate class A was correlated with the vigilance level. This suggests that duration and occurrence of microstates manifest different psychophysiological mechanisms. This view gets support from recent research that suggests that psychiatric disorders only affect one of these two microstates selectively (Michel & Koenig 2017). Importantly, to estimate the vigilance levels, we follow with the definition and algorithm described in (Falahpour et al 2018). Compared to alternative approaches like VIGALL (Hegerl & Hensch 2014, Olbrich et al 2009, Strauss et al 2015), the approach used here has the benefit that it produces continuous measures of vigilance and does not require electrooculography data.

### Role of Microstates A and B

We found negative associations between the occurrence of microstates A and B with the vigilance level. This observation goes in line with recent studies which link the presence of these two microstates to primary sensory processes. Microstate class B was suggested to be associated with the visual resting state network (Britz et al 2010, Custo et al 2017), a claim which was further supported by an increase in duration of the microstate class B in eyes-open rest (Seitzman et al 2017) and was shown to be associated with visual imagery thoughts (Lehmann et al 1998). The sources of the microstate class A seem to be in the temporal areas and can be associated with the auditory resting state network (Britz et al 2010, Custo et al 2017). While it does not affect our interpretation, it is noteworthy that a recent study (Milz et al 2016), however, contrary to the studies above, reported evidence that microstate class A could be related to visual and microstate class B to verbalization processes.

### Role of Microstate C

The functional role of the microstate class C is still unclear (Michel & Koenig 2017). Recent studies suggest a relation of microstate class C to cognitive control processes (Britz et al 2010). However, the decrease of the duration of microstate class C in serial subtraction tasks (Bréchet et al 2019, Seitzman et al 2017) and visualization tasks (Milz et al 2016) puts forward the hypothesis that it might reflect task-negative network activity (Michel & Koenig 2017). Our observation of a positive association of the duration of the microstate class C with vigilance favors the role of this microstate in cognitive control processes. Also, microstate class C is characterized by frontal to occipital topography with posterior predominance of activity. Taken together with the fact that induction of the different microstates is mainly determined by strength of the power in the alpha band (Milz et al 2017), an increased contribution of the microstate class C may reflect higher occipital alpha power. Loss of vigilance is characterized by a gradual shift of alpha power from the occipital towards frontal brain regions followed by a decrease in power in the alpha band and an increase in power in delta and theta frequency ranges that characterize drowsiness states (Olbrich et al 2009). Thereby the highest occipital alpha power is typically associated with the most vigilant state. Thus, it is not surprising, that we observe strong positive associations of the parameters of this microstate with vigilance.

### Role of microstate D

We observed a positive association between the duration of the microstate class D and the vigilance level. This could be explained by a number of recent studies suggesting that microstate class D is characterized by sources in middle and superior frontal areas as well as superior and inferior parietal areas (Britz et al 2010, Custo et al 2017) and has been hypothesized to be associated with the dorsal attention network. This hypothesis is further supported by the observation that occurrence and duration of this microstate increased during serial subtraction task (Bréchet et al 2019, Seitzman et al 2017). We note however that, somewhat contrary to this hypothesis, (Milz et al 2016) reported the highest duration and occurrence of the microstate class D during rest in comparison with a number of cognitive tasks. This suggests that microstate class D might reflect focus switching and reflexive aspects of attention. While we cannot reject or confirm any of these two based on these results, we find it noteworthy that both theories indirectly implicate the relation between microstate D and the vigilance state.

### Relation to the fMRI global signal

In line with recent studies (Falahpour et al 2018, Wong et al 2016, Wong et al 2013) we found a negative association between vigilance and the fMRI global signal. We additionally found that fMRI global signal was associated with the duration and contribution of microstate class C and the contribution of microstate class A. However, parameters of microstate class B as well as transition probabilities between microstate classes did not show any association with the global signal. This suggests that the change in vigilance may likely affect brain dynamics on multiple levels where the mechanisms that affect microstate parameters and global signal are essentially different.

### The potential mediating role of cingulate cortex

It is interesting to note that an increasing number of recent studies, using source reconstruction techniques, suggest that the cingulate cortex is a common source for all microstate classes (Custo et al 2017, Pascual-Marqui et al 2014). In line with these findings, multimodal imaging EEG-fMRI studies provided evidence that attenuation of vigilance levels leads to an increase in activity of the anterior part (Olbrich et al 2009) while caffeine intake leads to alterations of BOLD activity in the posterior part of the cingulate cortex (Falahpour et al 2018). Taken together, considering the role of cingulate cortex and arousal, these studies suggest the activity of the cingulate cortex to mediate the causal relation between vigilance level and appearance of microstates sequences.

### Prediction accuracy

We found that the observed association between parameters of the microstates and vigilance time series has predictive power. Using support vector machine regression, we could predict vigilance fluctuations with accuracy which is statistically significant but not numerically high. This suggests that vigilance levels and EEG microstate parameters are not reflecting necessary the same processes. Such view receives further support from observations that next to the power in the alpha frequency band, microstates are additionally related to delta, theta, and higher frequency bands (Khanna et al 2015). Taken together, and considering that vigilance level is a more fundamental characteristic of the organism, which is also partially modulated by the body, we speculate that the time course of microstates is influenced by vigilance through multiple systems.

### Analysis of microstates in psychiatric disorders

The finding that parameters of EEG microstates have temporal dynamics which are partly modulated by vigilance state has far reaching implications when microstates are studied in the context of psychiatric disorders. Patients suffering from psychiatric disorders often show abnormal sleep behavior and/or altered temporal vigilance structure. In light of the findings presented in this paper, the potential changes found in microstates could be essentially related to changes in vigilance which might or might not be a symptom or manifestation of the neural mechanism of the disorder, but indicate a rather straightforward change in sleep behavior.

### 4. Conclusion

We provided evidence for correlations and, crucially, causal relations between the fluctuations of vigilance level and temporal dynamics of the EEG microstates within the first 10 minutes of rest. We found that duration and contribution of microstate class C were positively, while occurrence and contribution of microstate classes A and B were negatively associated with vigilance. Changes in vigilance caused changes in EEG microstate parameters. The observed findings highlight the importance of taking vigilance levels into consideration in EEG microstate parameter investigations. We suggest that the cingulate cortex may be a potential mediator of the observations we made and the fact that EEG microstates reacted to the changes in vigilance level potentially by integrating multiple neural mechanisms.

## 5. Acknowledgements

MW was supported by DFG grant (Wa2674/4-10) and SFB779-A06. HJ was supported by fortüne grant of Medical Faculty of University of Tübingen (No. 2487-1-0). Data set 1 was collected in a trial sponsored by Biologische Heilmittel HEEL GmbH, Germany (NCT02602275) in which MW was a PI. EEG and fMRI data were provided as courtesy for the purpose of these analyses which were not related to the trial objectives. Data set 2 was collected in a study sponsored by German Research Foundation (DFG, WA 2673/10-1). Authors thank Prof. Andreas J. Fallgatter and Dr. Meng Li for the valuable feedbacks and support of the study. The authors declare no conflict of interest.

## 6. Data and code availability

The data of the dataset 2 and/or code used in the current study are available from the corresponding authors upon reasonable request subject to a formal data sharing agreement with Prof. Martin Walter. Dataset 1 can only be shared with formal agreement with Prof. Martin Walter and Biologische Heilmittel HEEL GmbH, Germany.

## 8. Appendix A: Association of the mean vigilance level and mean values of the microstate parameters

**Table A.1:** This table lists the results (r-values and p-values) of all preformed correlations between the mean vigilance level and mean values of the microstate parameters. Significant results (p < 0.05) are marked in bold. Mean duration and contribution of the microstate class C are positively, while occurrence and contribution of the microstates class A and B are negatively associated with vigilance level in both investigated data sets.

|  |  | A | B | C | D |
| --- | --- | --- | --- | --- | --- |
| Data set 1 | Duration | $r = -0.10$ $p = 0.579$ | $r = -0.02$ $p = 0.907$ | **$r = 0.61$ $p = 2 \times 10^{-4}$** | **$r = 0.46$ $p = 0.007$** |
| | Occurrence | **$r = -0.60$ $p = 2 \times 10^{-4}$** | **$r = -0.44$ $p = 0.009$** | $r = 0.17$ $p = 0.342$ | $r = -0.10$ $p = 0.566$ |
| | Contribution | **$r = -0.40$ $p = 0.019$** | **$r = -0.35$ $p = 0.046$** | **$r = 0.65$ $p = 4 \times 10^{-5}$** | $r = 0.07$ $p = 0.686$ |
| Data set 2 | Duration | $r = -0.19$ $p = 0.440$ | $r = -0.25$ $p = 0.308$ | **$r = 0.75$ $p = 2 \times 10^{-4}$** | $r = 0.37$ $p = 0.115$ |
| | Occurrence | **$r = -0.48$ $p = 0.037$** | **$r = -0.57$ $p = 0.010$** | $r = 0.45$ $p = 0.055$ | $r = -0.09$ $p = 0.699$ |
| | Contribution | $r = -0.41$ $p = 0.083$ | **$r = -0.48$ $p = 0.040$** | **$r = 0.67$ $p = 0.002$** | $r = 0.08$ $p = 0.731$ |

**Table A.2:** This table lists the results (r-values and p-values) of all preformed correlations between the mean vigilance level and mean values of the transition probability for transitions between four microstate classes. Significant results (p < 0.05) are marked in bold. For both data sets mean transition probabilities towards microstate class C are positively, while transition probabilities from microstate class D towards microstate class A is negatively associated with vigilance levels.

|  | Data set 1 | Data set 2 |
| --- | --- | --- |
| **A → B** | $r = -0.25$ $p = 0.150$ | $r = -0.26$ $p = 0.285$ |
| **A → C** | **$r = 0.65$ $p = 4 \times 10^{-5}$** | **$r = 0.61$ $p = 0.006$** |
| **A → D** | $r = -0.25$ $p = 0.152$ | $r = -0.29$ $p = 0.227$ |
| **B → A** | $r = -0.18$ $p = 0.298$ | $r = -0.40$ $p = 0.094$ |
| **B → C** | $r = 0.27$ $p = 0.117$ | **$r = 0.50$ $p = 0.027$** |
| **B → D** | $r = 0.09$ $p = 0.617$ | $r = 0.25$ $p = 0.309$ |
| **C → A** | $r = -0.32$ $p = 0.061$ | $r = -0.18$ $p = 0.449$ |
| **C → B** | **$r = -0.37$ $p = 0.030$** | $r = -0.33$ $p = 0.173$ |
| **C → D** | $r = 0.13$ $p = 0.464$ | $r = -0.07$ $p = 0.779$ |
| **D → A** | **$r = -0.40$ $p = 0.018$** | **$r = -0.61$ $p = 0.005$** |
| **D → B** | $r = -0.20$ $p = 0.264$ | $r = 0.04$ $p = 0.863$ |
| **D → C** | **$r = 0.48$ $p = 0.004$** | **$r = 0.48$ $p = 0.038$** |

## 9. Appendix B: Association of the vigilance time course and the time courses of the microstate parameters

**Table B.1:**
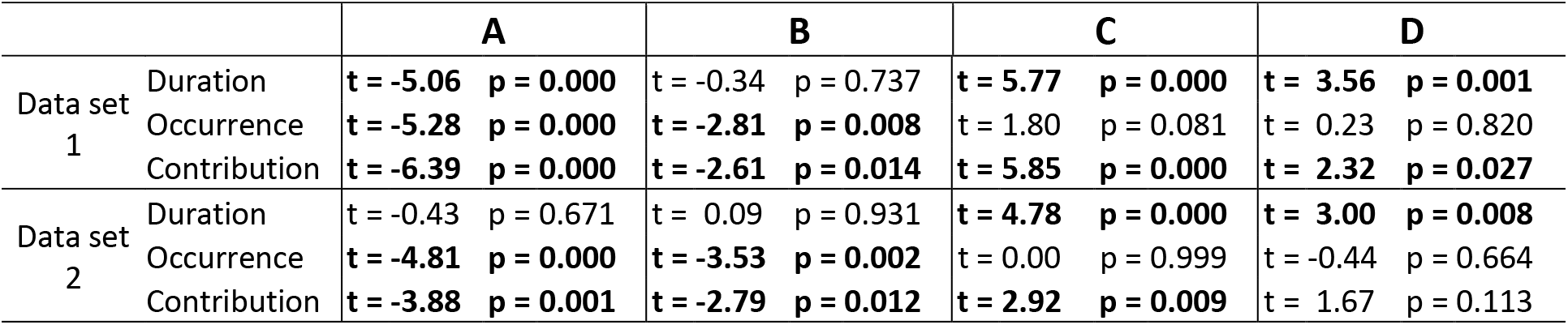
This table lists the results (t-values and p-values) of the one-sample t-test on the Fisher z-transformed subject-level correlation coefficients for the correlation the vigilance time-series and the time courses of the microstate parameters. Significant results (p < 0.05) are marked in bold. Time courses of the duration and contribution of the microstate class C as well as duration of the microstate class D are positively, while time courses of the occurrence and contribution of the microstates classes A and B are negatively associated with changes of vigilance level in both investigated data sets.

**Table B.2:**
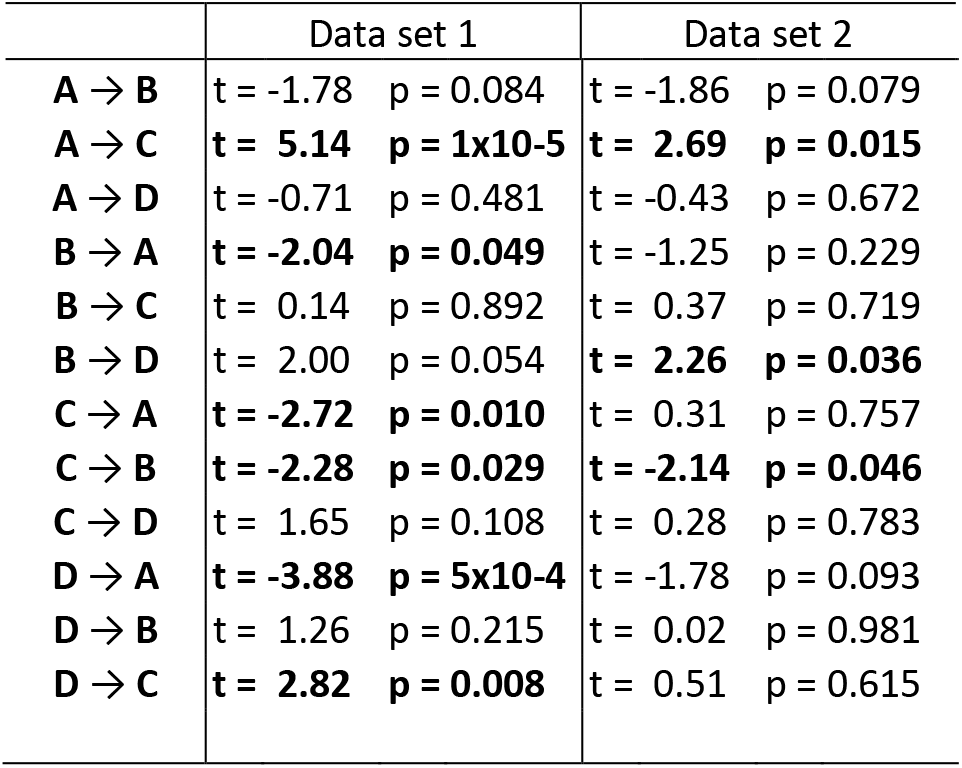
This table lists the results (t-values and p-values) of the one-sample t-test on the Fisher z-transformed subject-level correlation coefficients for the correlation the vigilance time-series and the time courses of the transition probabilities for transitions between four microstate classes. Significant results (p < 0.05) are marked in bold. Time courses of the transition probabilities for transitions from microstate class A towards microstate class C as well as transitions from for transitions from microstate class B towards microstate class D are positively, while time course for of the probability of transitions from microstate class C towards microstate class B is negatively associated with vigilance time-series.

## 10. Appendix C: Association of the global signal time-series and the time courses of the microstate parameters

**Table C.1:** This table lists the results (t-values and p-values) of the one-sample t-test on the Fisher z-transformed subject-level correlation coefficients for the correlation the global signal (Leonardi et al) time-series and the time courses of the microstate parameters. Significant results (p < 0.05) are marked in bold. Time courses of the duration and contribution of the microstate class C are negatively, while time course of the occurrence of the microstate class A is positively associated with GS time-series in both investigated data sets.

|  | A | B | C | D |
| --- | --- | --- | --- | --- |
| Data set 1 |  |  |  |  |
| Duration | t = 1.95 p = 0.060 | t = 1.50 p = 0.142 | **t = -2.77 p = 0.009** | t = -0.75 p = 0.456 |
| Occurrence | t = 1.70 p = 0.099 | t = 0.81 p = 0.423 | t = -1.07 p = 0.292 | t = -1.91 p = 0.065 |
| Contribution | **t = 2.24 p = 0.032** | t = 1.64 p = 0.110 | **t = -2.24 p = 0.032** | t = -1.43 p = 0.162 |
| Data set 2 |  |  |  |  |
| Duration | t = 2.08 p = 0.052 | t = 0.42 p = 0.680 | **t = -2.71 p = 0.014** | t = 0.00 p = 0.999 |
| Occurrence | t = 1.44 p = 0.168 | t = 0.38 p = 0.710 | t = -1.91 p = 0.072 | t = 0.90 p = 0.379 |
| Contribution | **t = 3.04 p = 0.007** | t = 0.81 p = 0.430 | **t = -2.64 p = 0.017** | t = 0.79 p = 0.441 |

**Table C.2:**
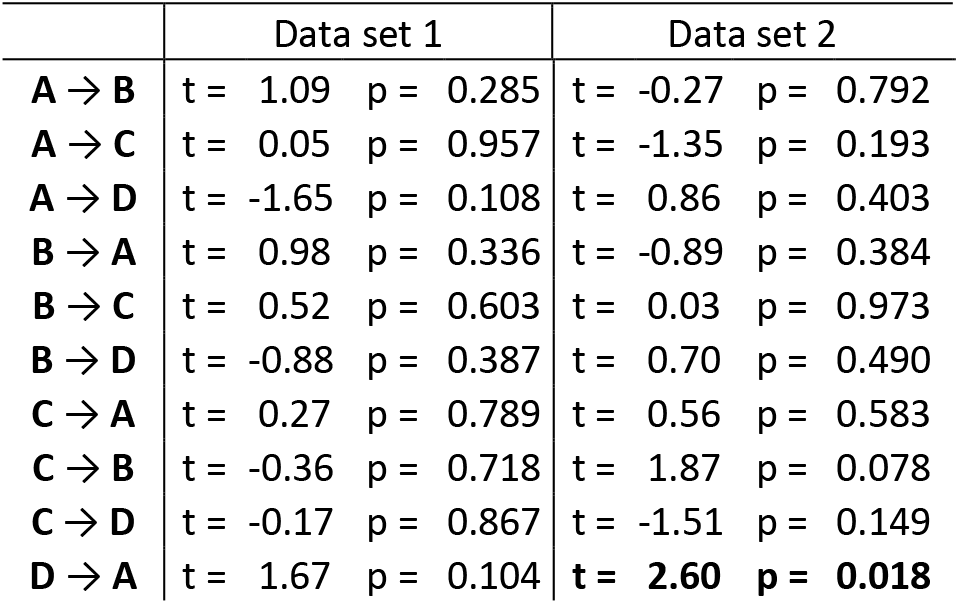
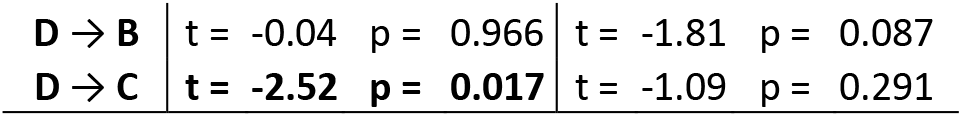
This table lists the results (t-values and p-values) of the one-sample t-test on the Fisher z-transformed subject-level correlation coefficients for the correlation global signal (Leonardi et al) time-series and the time courses of the transition probabilities for transitions between four microstate classes. Significant results (p < 0.05) are marked in bold. Association between global signal time course and time course of transition probabilities of microstates were weak and inconsistent across two investigated data sets.

